# Sex-specific responses to cold in a very cold-tolerant, northern *Drosophila species*

**DOI:** 10.1101/2020.07.26.221788

**Authors:** Darren J. Parker, Tapio Envall, Michael G. Ritchie, Maaria Kankare

## Abstract

Organisms can plastically alter resource allocation in response to changing environmental factors. For example, in harsh conditions organisms are expected to shift investment from reproduction towards survival, however, the factors and mechanisms that govern the magnitude of such shifts are relatively poorly studied. Here we compared the impact of cold on males and females of the highly cold-tolerant species *Drosophila montana* at the phenotypic and transcriptomic levels. Although both sexes showed similar changes in cold tolerance and gene expression in response to cold treatment, indicating that the majority of changes are concordant between the sexes, we identified a clear reduction in sexually dimorphic gene expression, suggesting that preparing for colder season also involves reducing investment in sex-specific traits. This reduction was larger in males than females, as expected if male sexual traits are more condition-dependent than female traits, as predicted by theory. Gene expression changes were primarily associated with shifts in metabolic profile which likely play a role in increasing cold tolerance. Finally, we found that the expression of immune genes was reduced following cold treatment, suggesting that reduced investment in immunity may be important in helping flies survive colder periods.

## Introduction

Life history strategies involve strategic allocation of investment between reproduction and survival, and relative investment in these depends on a wide range of intrinsic and extrinsic factors [1–3]. Many of these factors vary throughout an organism’s lifetime meaning selection will favour different allocations at different times. As a result, organisms are typically able to plastically shift the relative allocation of resources in response to environmental cues, particularly when changes in the environment are predictable [4,5].

One predictable shift is the change from summer to winter when temperature is decreasing and day length is shortening. For organisms at high latitudes, this harshening of the environment is expected to produce a shift in resource allocation with a greater investment into survival over reproduction. This is because, in order to survive the colder conditions, organisms need to produce a range of costly metabolites and proteins and to begin to store resources to survive the colder season [6,7]. Numerous factors are likely to influence the magnitude of these trade-offs including life cycle, age, and condition. One potentially important factor, which is however surprisingly rarely studied, is that of sex. With the changing of the seasons both males and females have to adjust to the same conditions, so we may expect that the physiological shifts to survive colder temperatures may be similar. However, the relative costs of coping with lower temperatures may differ between the sexes. For instance, males may be more susceptible to cold than females (e.g. in *D*. *melanogaster* [8]) meaning that a greater shift in resources would be required in order for males to survive colder periods. In addition, although sexual traits in both sexes are expected to be reduced in response to worsening conditions [9–12], it is also expected that condition-dependence will be stronger for males than females due to sexual selection [13,14], meaning we should expect a larger shift in resources in males than in females when they prepare for the onset of cold. Finally, males and females typically have very different expression profiles, expressing a large fraction of genes at different levels throughout the genome to produce sexually dimorphic phenotypes [15,16]. Such differences in expression could restrict additional changes in gene expression (for instance if increased expression would have negative effects in one sex but not the other [17] or if a gene is already maximally expressed in one sex but not the other) and thus produce differences in how each sex can respond to cold.

Here we have three objectives. Firstly, we examine if males and females have similar phenotypic responses to the onset of cold in a cold-adapted species, *Drosophila* (*D*.) *montana*. This species is particularly well adapted to cold environments [18,19] with both sexes able to overwinter as adults at high latitudes and altitudes meaning that being able to survive cold-stress is an important part of their life history. Secondly, we examine if males and females have similar changes in gene expression when subjected to cold using an RNA-seq approach. This approach is ideal for examining how shifts in resource allocation occur since these changes are plastic and thus differences in gene expression will reflect differences in resource allocation strategies. We predict that i) males and females will show similar phenotypic changes to cold, ii) both sexes will show similar changes in gene expression for most genes, iii) genes associated with producing sexual differences in traits (i.e. sex-biased genes) will show significant reductions in expression in response to cold and, iv) this reduction will be larger in males than in females. Finally, we examine the functional processes associated with genes that change expression to gain insight into the molecular mechanisms by which males and females cope with the onset of cold.

## Results

### Males and females show a similar phenotypic response to cold

To assess if males and females have a similar response to cold we experimentally reduced the temperature flies were maintained at from 19 °C to 6 °C, representing average daytime temperatures in Central Finland in late July and early October respectively (www.worlddata.info). After 5 days we compared the critical thermal minimum (CTmin, the temperature at which flies lose neuromuscular function) of cold treated flies to control flies (see methods for details). Both males and females showed a significant increase in cold tolerance i.e. a lower CTmin following cold treatment (Fig. 1, Table 1). There was no significant treatment by sex interaction indicating that males and females have a similar phenotypic shift in cold tolerance (Table 1).

**Table 1.** The effect of sex, cold treatment, and weight assessed by ANOVA (full model). Significant p-values are in bold.

| Model Term | Sum of Sq | Df | F values | P |
| --- | --- | --- | --- | --- |
| Sex | 0.0038 | 1 | 0.0391 | 0.8436 |
| Cold Treatment | 1.0815 | 1 | 11.1732 | **0.00108** |
| Weight | 0.0838 | 1 | 0.8657 | 0.35386 |
| Sex * Cold Treatment | 0.1345 | 1 | 1.39 | 0.24053 |
| Residuals | 12.7769 | 132 |  |  |

**Fig. 1.**
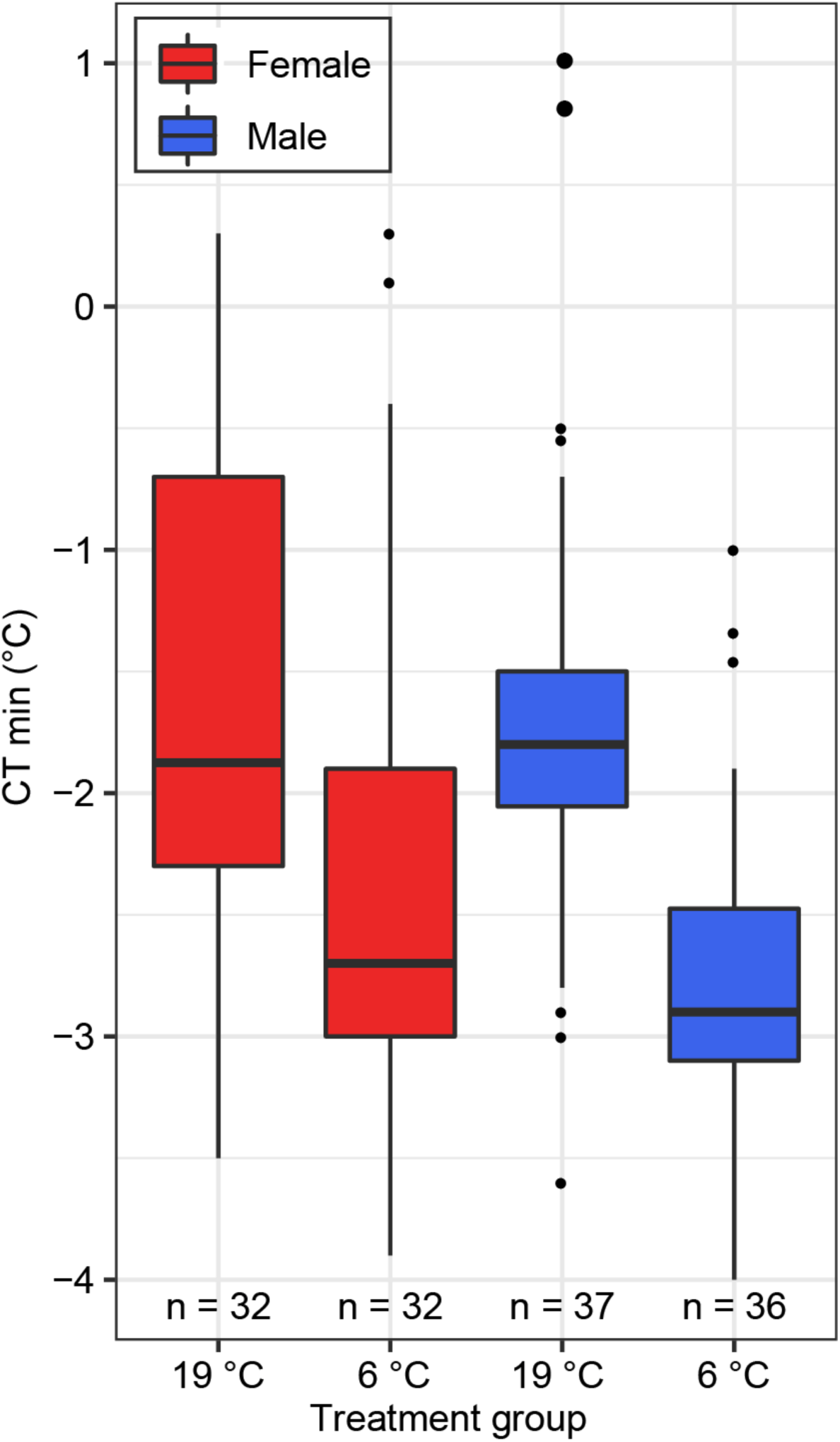
Cold tolerance is higher for male and females when maintained at a colder temperature. Treatment group indicates whether flies were maintained at 19 °C or 6 °C for five days (see text for detailed methods). CTmin is the temperature at which flies lose neuromuscular function.

### Males and females show similar changes in gene expression in response to cold

Flies raised under the same conditions used for phenotypic measurements (above) were also used for gene expression analyses. Both sex and temperature strongly influenced gene expression (Fig. 2A) with samples clustering first by sex, then by temperature (Fig. 2B). Differential expression analyses found that a little over 10% of all expressed genes were DE in response to cold in both males (1236 / 9338) and females (1062 / 9338), with significant overlap between genes DE in males and females (Fig. 3). This overlap was much greater than expected by chance (p = 1.7 × 10^−110^). Gene expression change in response to cold was also highly correlated between males and females for all genes (rho = 0.59, p-value < 2.2 × 10^−16^, Fig. 4), and for genes DE in either males or females (rho = 0.73, p-value < 2.2 × 10^−16^) with only a small number of genes (64) showing a significant sex by temperature interaction (Figs. 3, S1).

**Fig. 2.**
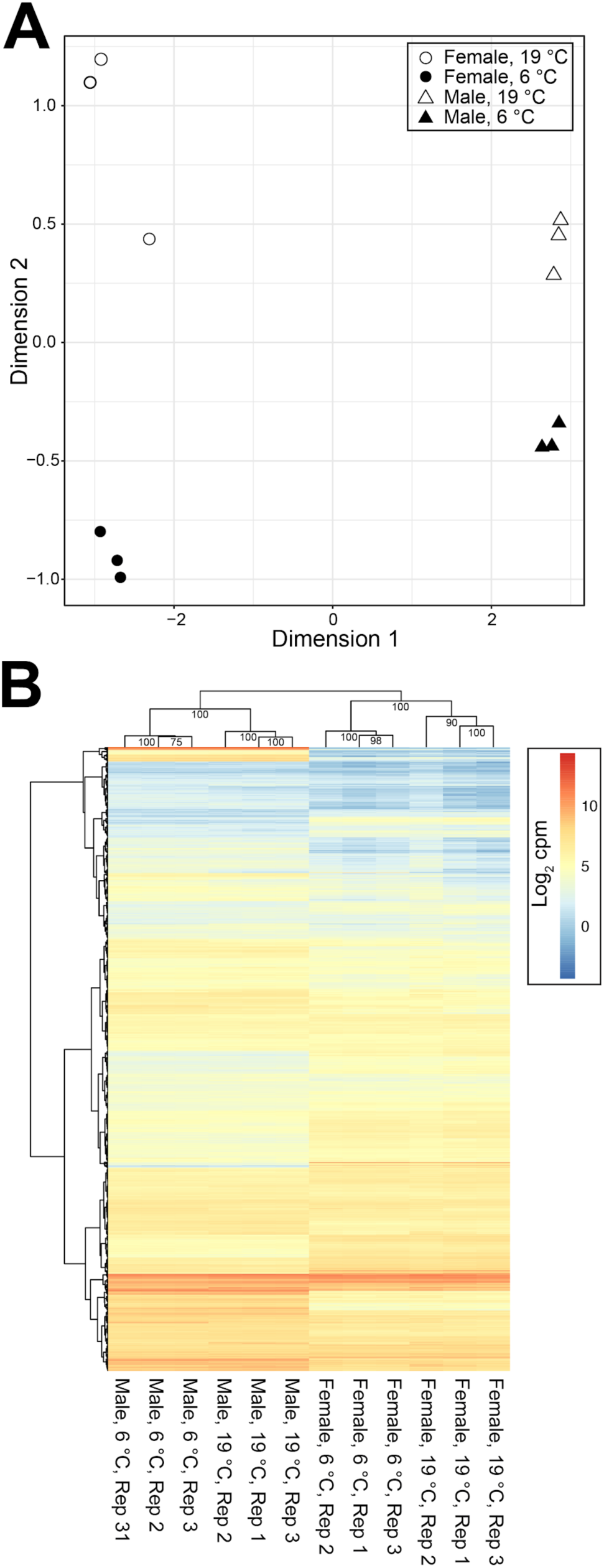
**A)** MDS plot of male (triangle) and female (circle) expression when kept at 19 °C (empty shapes) or 6 °C for 5 days (filled shapes). Distances between samples in the MDS plot approximate the log2 fold change of the 500 genes with the largest biological variation between the libraries. **B)** Heatmaps and hierarchical clustering of gene expression (log_2_ CPM). Values on each node show the bootstrap support from 1000 replicates.

**Fig. 3.**
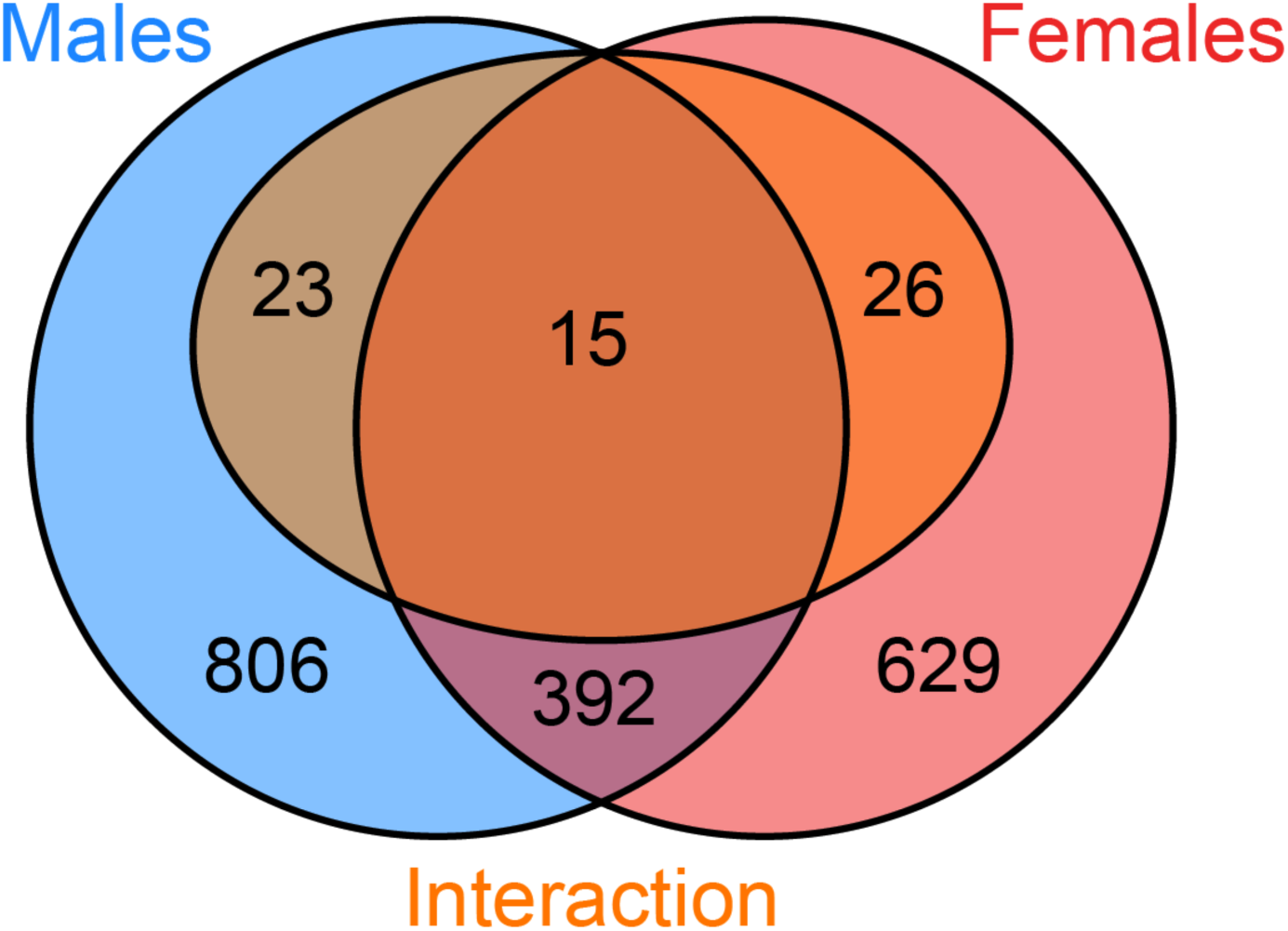
Venn-diagrams showing the overlap of genes DE in response to cold treatment in males (blue) and females (red). Genes showing a significant sex by treatment interaction are shown in orange.

**Fig. 4.**
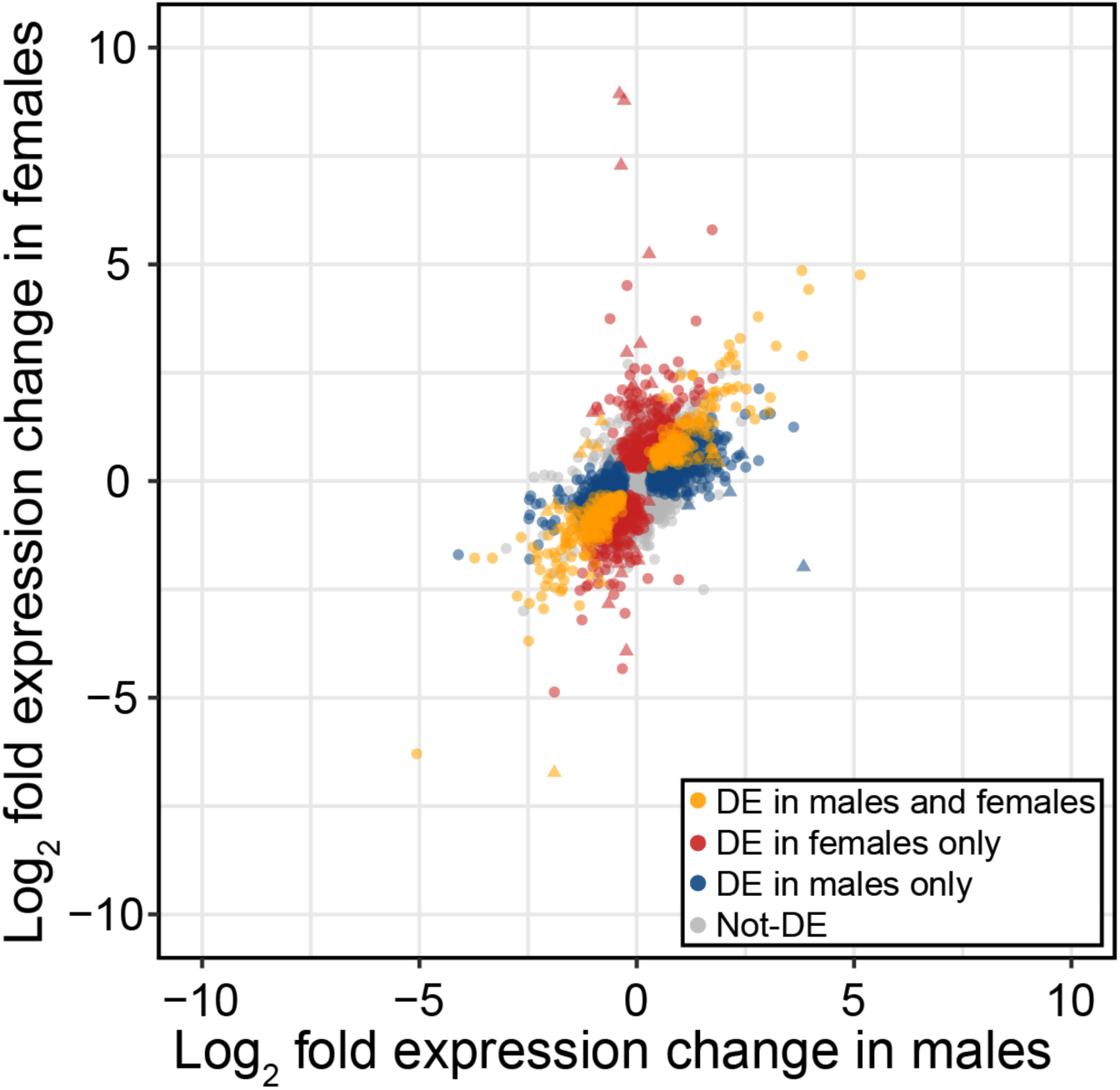
Expression change in males and females in response to cold treatment, indicating genes differentially expressed in males and females (orange), females only (red), males only (blue) and in neither sex (grey). Triangle points indicate a significant sex by treatment effect.

### Sexually dimorphic gene expression is reduced in response to cold

To determine if exposure to cold alters the amount of sexually dimorphic gene expression we first examined how sex-biased genes change in response to cold. We found the amount of sexually dimorphic gene expression decreased in both males and females, with male-biased genes reducing in expression and female-biased genes increasing in expression in males, and female-biased genes decreasing in expression in response to cold in females (Fig. 5). Shifts in sex-biased genes were larger in males than females, for both male- and female-biased genes, leading to a more ‘feminised’ transcriptome overall (Fig. 6). Note, similar shifts in sex-biased gene expression were also found when using a more conservative set of sex-biased genes (genes that are sex-biased in both *D*. *montana* and its close relative *D*. *virilis* (Fig. S2; S3)), however, changes in expression in females were no longer significant. Next, we examined the correlation of gene expression for males and females for all genes in each temperature treatment. Correlations between male and female gene expression were significantly higher for flies kept at 19 °C than those at 6 °C (Fisher’s z test, p = 0.0163), though the magnitude of this difference was small (19 °C *r* = 0.686, 6 °C *r* = 0.704).

**Fig. 5.**
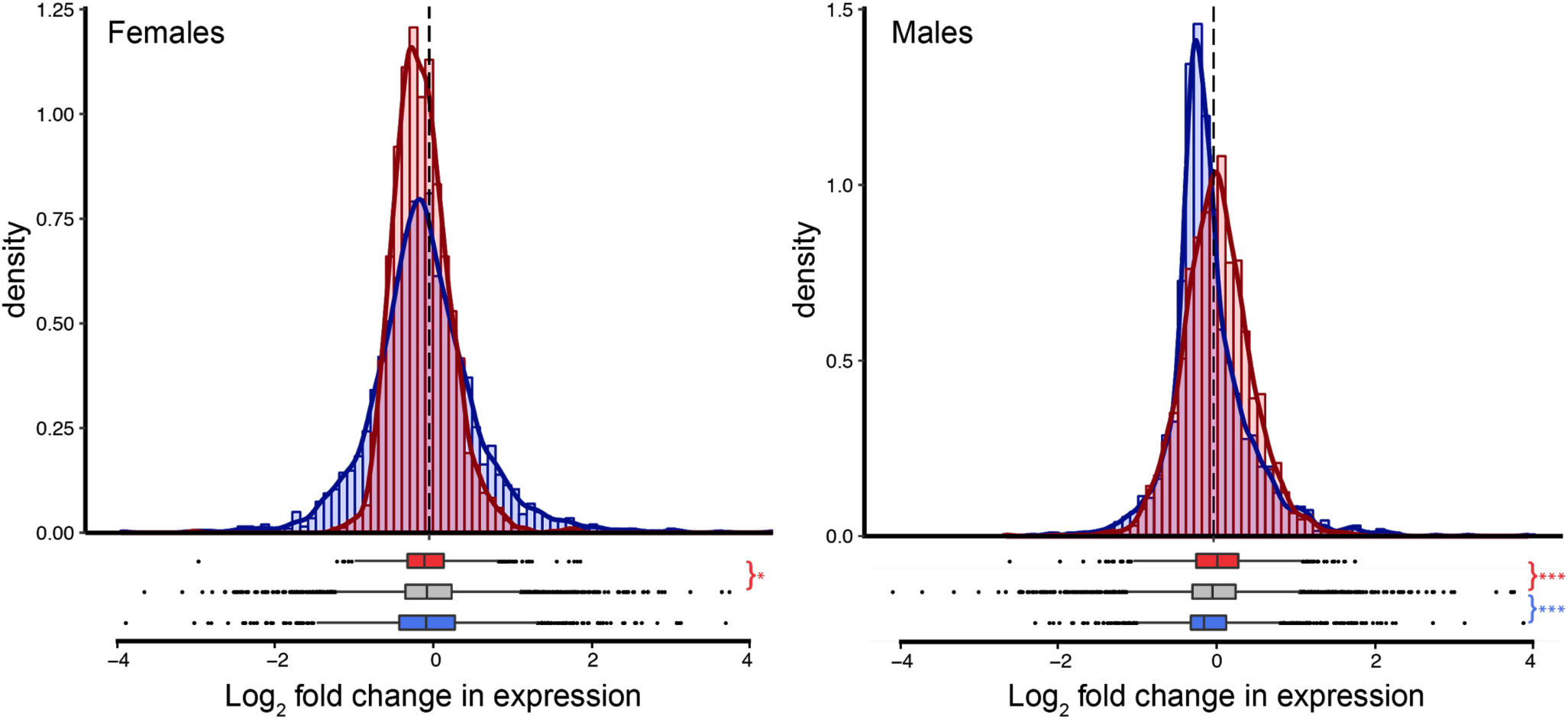
Expression shifts in sex-biased genes following cold treatment in females and males. Positive values indicate increased expression in cold-treated flies. Asterisks indicate the significance level (FDR) of Wilcoxon tests comparing the change in expression in female-biased (red) and male-biased (blue) genes to unbiased genes (***<0.001, **<0.01, *<0.05).

**Figure 6.**
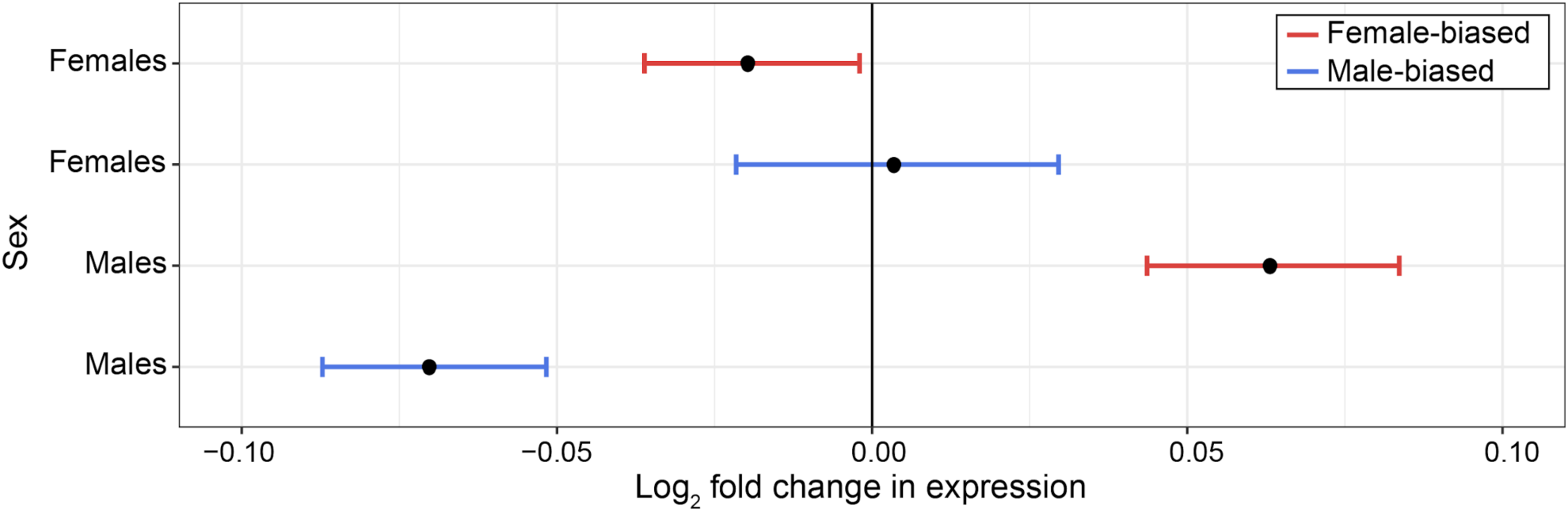
Expression shifts in male- (blue) and female- (red) biased genes following cold treatment in females and males relative to the median expression of unbiased genes. Positive values indicate increased expression in cold-treated flies. Points indicate pseudo-median and error bars indicate the 95% confidence interval.

### Genes differentially expressed in response to cold are enriched for metabolism and immune response in males and females

We used functional annotation clustering to examine the function of genes DE in response to cold. Specifically, we used this approach to identify processes enriched for genes DE in both males and females, genes DE in males only, genes DE in females only, and genes showing a sex by treatment interaction (Supplemental Table 1). The largest number of enriched gene clusters were found from genes DE in both sexes. These were primarily connected to metabolism (e.g. lipid metabolism, fatty acid biosynthesis, carbohydrate kinases, metalloproteases, and aminotransferases) as well as to the immune response (e.g. innate immune response and DM9 repeat) suggesting these processes are important for both males and females to adjust to a colder environment. Interestingly, all genes in immune-related clusters showed decreased expression following cold treatment (Fig. S4), suggesting a reduced investment in immune function. Genes showing a significant sex by treatment interaction were enriched for transmembrane transport, suggesting that while changes to this process are important for both males and females in a colder environment, how it is mediated differs between the sexes (Fig. S5). Finally, we found that the functional clusters enriched in genes DE only in females or only in males were different. In females, clusters were related to oxidoreductase activity and the biosynthesis of amino acids, whereas in males, clusters were related to cytoplasmic translation, protein biosynthesis, transmembrane transport, ATP-binding domain, glutamine metabolic processes, and nucleotide-binding.

## Discussion

In order to survive harsh conditions organisms are expected to shift investment towards survival and away from reproduction. This shift may differ between the sexes due to relative differences in the costs of reproduction or survival, or because of differences in regulatory architecture [15,16]. Despite its importance, the underlying mechanisms responsible for shifts in investment are poorly studied [20]. Here we examined such a shift by investigating the phenotypic and transcriptomic changes associated with the onset of cold in males and females of a cold tolerant fly species, *D*. *montana*.

We found that both sexes show a similar change in phenotype following cold treatment. Changes in cold tolerance (measured by CTmin) might be expected to be similar as both males and females have similar baseline cold tolerances in benign temperatures [21] and because both sexes need to adjust their physiology in order to survive in colder temperatures. Although cold tolerances are similar between the sexes at benign temperatures, this need not be the case. For instance, in the more temperate, but closely related species, *D*. *virilis*, males show much lower cold tolerances than females at benign temperatures [21]. Why this is not the case in *D*. *montana* is unclear, but one possibility is that *D*. *montana’s* ability to survive extremely cold temperatures constrains cold tolerance at warmer temperatures in both sexes.

Changes in gene expression in response to cold were similar between the sexes, suggesting that males and females adjust their physiology using largely the same mechanisms. Unfortunately, very few studies have examined gene expression changes in males and females in response to stressful environmental conditions, however, our results agree with a previous study showing that most genes in male and female *D*. *melanogaster* are concordantly regulated in response to changes in dietary composition [23]. While we found that most gene expression differences were similar between the sexes, it is clear that there are also differences. We observed a reduction in sexually dimorphic gene expression, with males and females having more similar expression in the harsher, colder condition. This finding agrees with previous studies in *D*. *melanogaster* [24] and beetles [25–27] which showed a reduction of sexually dimorphic gene expression with reduced environmental quality. The shifts in sex-biased gene expression we observed were relatively small overall, but it is notable that the shifts were larger in males than females. This is likely because investment into sexual traits is more condition-dependent in males than females [13,14], suggesting that a reduction of investment into expensive male functions during winter represents a greater change in life history than female changes (see also [24,28]). Relatively few genes showed a significant sex by treatment interaction, reinforcing the idea that most changes in gene expression are concordant between the sexes. Interestingly however, these few genes were enriched for transmembrane transport, a process that has been previously associated with cold adaptation in a number of insects [29], including *D*. *montana* females [30]. Changes to transmembrane transport are thought to be particularly important for surviving cold temperatures by preventing a loss of cellular ionic balance [31–34]. Our results suggest that, despite its importance, changes to transmembrane transport are mediated by different genes in each of the sexes. The reasons for this are not clear, but it is possible that these differences may arise from sex-specific genetic constraints.

Overall, we found that the transcriptomic response to cold is extensive, with several hundred genes showing differential expression. By examining the functional processes associated with these gene expression changes we are able to gain insights into the mechanisms by which males and females cope with the onset of cold. We did this in two separate analyses, first examining processes enriched in genes DE in both sexes, then processes enriched in genes DE only in male or only in females. Processes enriched in both sexes included many that have been previously associated with increasing cold tolerance including metabolic shifts in lipids and carbohydrates. By altering their metabolic profile insects are able to maintain osmotic balance and stabilize the membrane structures of a cell as temperatures decrease (e.g. [35– 38]. In particular, changes in lipids, fatty acids, and polyols have been shown to be important for cold adaptation in many insect taxa [6,7,39,40], including *D*. *montana* females [19,30,41]. This is consistent with the changes we identify here, including DE of previously identified candidate genes *Inos* and *CG6910* [30,41]. Both of these genes belong to the inositol biosynthetic pathway, emphasizing the importance of this pathway for surviving colder temperatures in *D*. *montana*.

Interestingly, we also found an enrichment of immune-related processes including innate immune function and genes with a DM9 repeat (which likely have an antimicrobial function [42,43]). Genes enriched for these processes reduced in expression following cold treatment, suggesting that investment in immune function is reduced in colder temperatures. This finding is in contrast to most previous work which shows that cold exposure stimulates an increase in immune function (e.g. *Ostrinia furnacalis* [44], *Pyrrharctia isabella* [45], *Megachile rotundata* [46], and *D*. *melanogaster* [47–49]). Such increases may represent a shift in investment for enhanced immune activity during colder periods, however, it is also consistent with a general stress response, or immune activation due cold-induced tissue damage [50]. Since we observe a reduction in immune expression, the changes we see in immunity are unlikely to be as a result of general stress or tissue damage response but instead due to a specific reduced investment in immunity. The only other study to our knowledge that found a reduction in immune function due to cold was performed with *Gryllus veletis* [51], which like *D*. *montana* has an overwintering diapause stage (though as a nymph rather than an adult). Unlike other studies that used insects with an overwintering diapause stage (e.g. *Megachile rotundata* [46]) both our and Ferguson et al.’s [51] study also used the developmental stage that will eventually enter into diapause for experiments. As such the cold treatment used in our and Ferguson et al.’s [51] study mimics the changing of the seasons and thus may cue these insects into preparing for the onset of winter. In these cases, reducing resource allocation in immunity is likely to be beneficial as maintaining immune function is energetically costly [52] and insects need to conserve energy reserves to survive the winter [53,54]. As such, reducing investment in immune function may be a common adaptation for insects preparing to overwinter, but future work in other species will be required to determine if this is a general phenomenon.

Processes enriched in genes DE in only one of the sexes are diverse. These processes represent changes that are potentially more important for one sex than the other, however, since few genes showed a significant sex by treatment interaction these processes are also likely to have some role in both sexes. Although diverse, most of the enriched processes are involved in metabolism, including: biosynthesis of amino acids and proteins, and glutamine metabolic processes, all of which have been previously associated with increased cold tolerance in insects [7,55–58]. In addition, we also found an enrichment of processes associated with oxidoreductase activity in females, which may help flies defend against increased oxidative stress induced by exposure to cold [47].

In conclusion, we found that male and females respond to the onset of cold in a similar way at both the phenotypic and transcriptomic levels. Despite this, cold treatment also reduced the amount of sexually dimorphic gene expression, with males showing a larger reduction than females, suggesting that preparing for colder periods involves reducing investment in male-specific functions. Gene expression changes were mainly associated with shifts in metabolic processes, however, we also observed decreased expression of immune genes suggesting that reduced investment in immunity may be an important adaptation to help survive the colder season. Finally, our results suggest that sex-specific adaptations involved in life history tradeoffs are subtle but potentially important, even when they are not apparent at the phenotypic level, highlighting the importance of examining tradeoffs at both phenotypic and molecular levels.

## Methods

### Samples

A genetically variable population cage was established using twenty fertilized *D*. *montana* females collected in 2013 from Korpilahti (62°N), Finland. This population cage was maintained in constant light at 19 °C to prevent the flies from entering diapause but note that *D*. *montana* do not lose circadian clock rhythmicity in constant light in contrast to *D*. *melanogaster* [59]. Newly enclosed flies from the cage population were anaesthetized with CO_2_ and separated by sex under a microscope and placed into half-pint bottles with yeast-malt medium. For the next 16 days, the bottles were kept at 19 °C, and flies were transferred to new bottles every week. After 16 days, half of both females and males were subjected to the cold treatment, which was 5 days at 6 °C [19] and the rest of the flies served as a control group remaining at 19 °C. At 21 days, we performed phenotyping and RNA extractions. Note that different individuals were used for phenotyping and RNA-extractions.

### Phenotypic measurements of cold tolerance

The phenotypic effect of cold on cold tolerance was determined by measuring the critical thermal minimum (CTmin) of the flies. CTmin is the temperature at which flies lose neuromuscular function, causing them to fall into a reversibly immobilized state called chill-coma (see Andersen et al. [60] for details). 21-day-old flies were put into 10 cm long glass tubes of diameter 1 cm, with 2-3 flies per tube. Flies in tubes were kept apart by pieces of plastic foam. The tubes were then sealed with Parafilm and submerged into a 30 % glycol-water mixture within a Julabo F32-HL refrigerated/heating circulator. Temperature of the liquid was then decreased at the rate of 0.5 °C/min from 19 °C to −10 °C as to be slow enough to allow the insect’s body temperature to cool with the temperature in the chamber but fast enough to avoid a substantial physiological response, during the cooling [61].

CTmin was recorded as the temperature at which a fly entered a chill-coma state by falling down. The experiment was done in batches of no more than 8 tubes with a maximum of 24 flies for a total of 137 flies. Flies were cooled to a point at least a couple of degrees Celsius below the temperature at which the last fly had entered chill-coma. Afterwards, the tubes were incubated at room temperature until all the flies had recovered from the chill-coma, to make sure that all the flies were normal, healthy individuals. Finally, the flies were killed by putting them into a freezer (−20 °C) for at least 12 hours, after which their weight was measured. The effects of temperature, sex, temperature by sex interaction, and weight on (log-transformed) critical thermal minimum were then tested using a type-III ANOVA in R (v. 3.5.1) [62]. Note that a value of five was added to all values of CTmin before log-transforming data to avoid taking the log of negative values.

### RNA extraction and sequencing

21-day-old flies were collected from the maintenance chambers and flash-frozen in liquid nitrogen and pooled into 12 samples with three flies in each sample for both cold treated and control groups. Flies were crushed with a plastic mortar, after which RNA extraction using ZR Insect & Tissue RNA Micro Kit with DNase treatment (Zymo Research) was carried out. RNA concentration was measured with Qubit (ThermoFisher), purity with NanoDrop ND-1000 (NanoDrop Technologies) and integrity with TapeStation 2200 (Agilent Technologies). Strand-specific library preparation (one library per sample) and paired-end (150 + 150 bp) Illumina sequencing (Illumina HiSeq 3000, 5 lanes) was then performed at the Finnish Functional Genomics Center (FFCG), Turku, Finland.

### Read trimming and mapping

Raw reads were trimmed before mapping. Firstly, CutAdapt [63] was used to trim adapter sequences from the reads before further trimming reads using Trimmomatic v 0.36 [64]. All reads were trimmed to 140 bp, then quality trimmed with the following options: LEADING:30 TRAILING:30 SLIDINGWINDOW:17:19. Any reads less than 85 bp in length after trimming were discarded. Quality-trimmed reads from each library were then mapped separately to the *D*. *montana* reference genome [65] using STAR (v. 2.4.2a) [66] with default options. Read counts for each gene were then obtained using HTSeq (v. 0.9.1.) [67].

### Differential gene expression analysis

Expression analyses were performed using the Bioconductor package EdgeR (v. 3.24.0) [68] in R (v. 3.5.1) [62]. Genes with counts per million <0.5 in 2 or more libraries per condition were excluded. Normalization factors for each library were computed using the TMM method. To estimate dispersion, we fit a generalized linear model (GLM) with a negative binomial distribution with the terms sex, temperature and their interaction. A quasi-F test was used to determine the significance of model terms for each gene by comparing appropriate model contrasts, with p-values corrected for multiple tests using Benjamini and Hochberg’s algorithm [69]. Statistical significance was set to 5%. Whether genes DE in males and females showed a greater overlap than expected by chance was determined using the SuperExactTest package (v. 0.99.4) [70].

Sex-biased genes were classified as genes showing significant differences (FDR <0.05, |log2FC| ≥ 1) between males and females in both control and cold treated samples. We chose these thresholds in order to select a robust set of sex-biased genes and to reduce the effect of sex-biased allometry [71]. Changes in expression of sex-biased genes in response to cold were then determined using a Wilcoxon test, corrected for multiple tests using Benjamini and Hochberg’s algorithm. We also repeated this analysis when sex-biased genes were defined with the additional condition that they are also sex-biased in *D*. *virilis*. Values for sex-biased expression in *D*. *virilis* were obtained from the sex-associated gene database [72] (downloaded 5^th^ November 2019). Genes sex-biased in *D*. *montana* and *D*. *virilis* showed good agreement (Fig. S3). To examine the overall similarity of male and female gene expression in each condition, we compared Spearman’s correlation coefficients of male and female gene expression (as mean log_2_ CPM) in control and cold treated flies using a Fisher’s z test implemented in the cocor package [73] in R [62].

### Functional annotation clustering

Functional annotation clustering of DE genes was carried out using DAVID (Database for Annotation, Visualization, and Integrated Discovery) v. 6.8 [74,75] with *D*. *melanogaster* orthologs (obtained from www.flymine.org). When multiple orthologs were obtained one was chosen at random to be used in DAVID. DAVID clusters genes into functional groups using a “fuzzy” clustering algorithm, and then uses a Fisher’s exact test to identify significantly enriched functional groups. A functional group was considered to be significantly enriched if its enrichment score was greater than 1.3 (p < 0.05).

## Data accessibility

Raw reads have been deposited in SRA under accession codes SRR10960337 - SRR10960348 (see Supplemental Table 2). Scripts for the analyses in this paper are available at https://github.com/DarrenJParker/montana_sex-specific_responses_to_cold and will be archived at Zenodo after acceptance. Raw CTmin data is given in Supplemental Table 3. Full gene expression results are given in Supplemental Table 4.

## Acknowledgements

This work was supported by Academy of Finland projects 268214 and 322980 to MK and a NERC (UK) grant NE/P000592/1 to MGR.

## Author Contributions

M.K. and D.J.P. designed the study. M.K. and T.E. collected samples and performed molecular work. D.J.P. and T.E analysed the data with input from M.G.R and M.K. D.J.P., M.G.R and M.K. wrote the manuscript with input from T.E.

