## Supplementary material for "Sex-specific responses to cold in a very cold-tolerant, northern *Drosophila species*": Supp Figs and Tables 1-2

**Supplemental Table 1 | Enriched functional processes**

| Name | p | Sex | Description |
|---|---|---|---|
| B1 | 0.002 | Both sexes | Lipid metabolism, Fatty acid biosynthesis, Fatty acyl-CoA synthase |
| B2 | 0.003 | Both sexes | DM9 repeat |
| B3 | 0.005 | Both sexes | Keto-sugar kinase, carbohydrate kinase |
| B4 | 0.010 | Both sexes | Flavin adenine dinucleotide binding, Flavoprotein |
| B5 | 0.011 | Both sexes | Carboxypeptidase, metalloprotease |
| B6 | 0.014 | Both sexes | Pyridoxal phosphate-dependent transferase |
| B7 | 0.014 | Both sexes | signal peptide, Secreted, disulfide bond |
| B8 | 0.022 | Both sexes | Carboxypeptidase, metallocarboxypeptidase activity |
| B9 | 0.031 | Both sexes | wax biosynthetic process, Fatty acyl-CoA reductase, long-chain fatty-acyl-CoA metabolic process |
| B10 | 0.040 | Both sexes | oxidoreductase activity, D-isomer specific 2-hydroxyacid dehydrogenase, NAD-binding, catalytic domain |
| B11 | 0.045 | Both sexes | nucleotide phosphate-binding region:NAD, active site:Proton acceptor |
| B12 | 0.049 | Both sexes | Innate immune response, immunity |
| F1 | 0.024 | Females only | Oxidoreductase, oxidoreductase activity, oxidation-reduction process |
| F2 | 0.024 | Females only | Glycolytic process, glycolysis, biosynthesis of amino acids |
| M1 | 0.000 | Males only | Cytoplasmic translation, structural constituent of ribosome, ribosomal protein |
| M2 | 0.011 | Males only | Protein biosynthesis, translational initiation |
| M3 | 0.022 | Males only | transmembrane transport, Major facilitator superfamily domain |
| M4 | 0.037 | Males only | DEAD-box, Helicase, ATP-binding domain |
| M5 | 0.038 | Males only | Glutamine metabolic process, glutamine amidotransferase |
| M6 | 0.048 | Males only | ATP binding, Nucleotide-binding |
| I1 | 0.027 | Interaction | Transmembrane transport, integral component of membrane |

**Supplemental Table 2 | Accession numbers for sequenced samples**

| Accession number | Temperature | Sex | Replicate number |
|---|---|---|---|
| SRR10960345 | 6 | female | 1 |
| SRR10960337 | 6 | female | 2 |
| SRR10960346 | 6 | female | 3 |
| SRR10960342 | 19 | female | 1 |
| SRR10960341 | 19 | female | 2 |
| SRR10960343 | 19 | female | 3 |
| SRR10960339 | 6 | male | 1 |
| SRR10960340 | 6 | male | 2 |
| SRR10960338 | 6 | male | 3 |
| SRR10960348 | 19 | male | 1 |
| SRR10960347 | 19 | male | 2 |
| SRR10960344 | 19 | male | 3 |

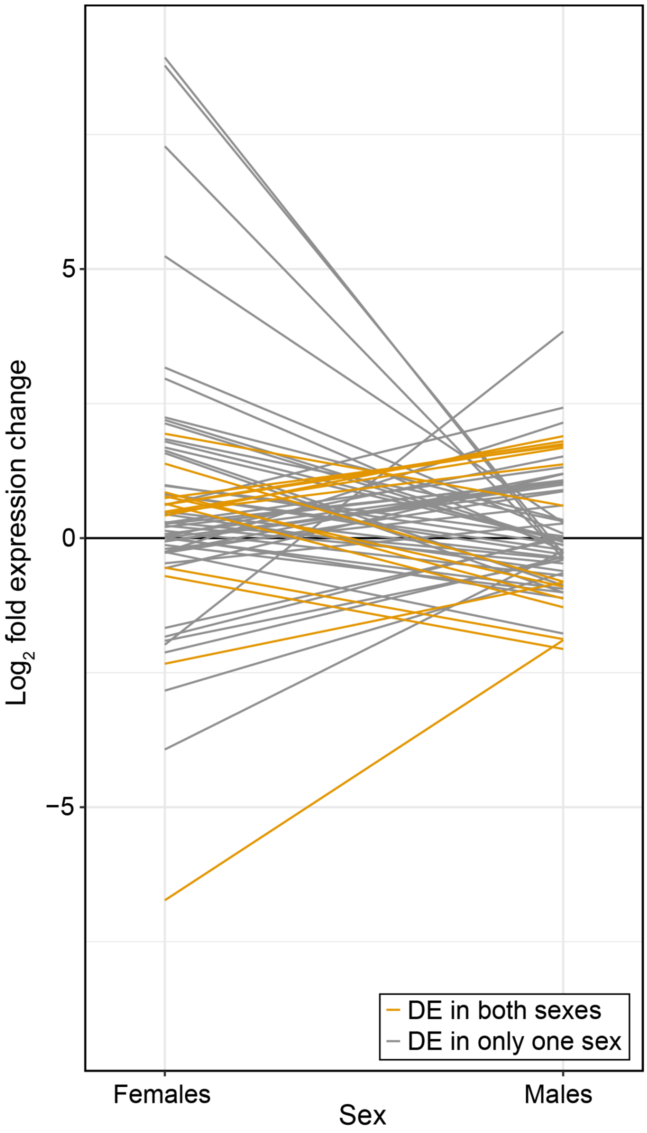


**Fig. S1 |** Expression change in males and females for genes with a significant sex by treatment interaction**.** Orange = genes DE in both males and females. Grey = genes DE in either males or females.

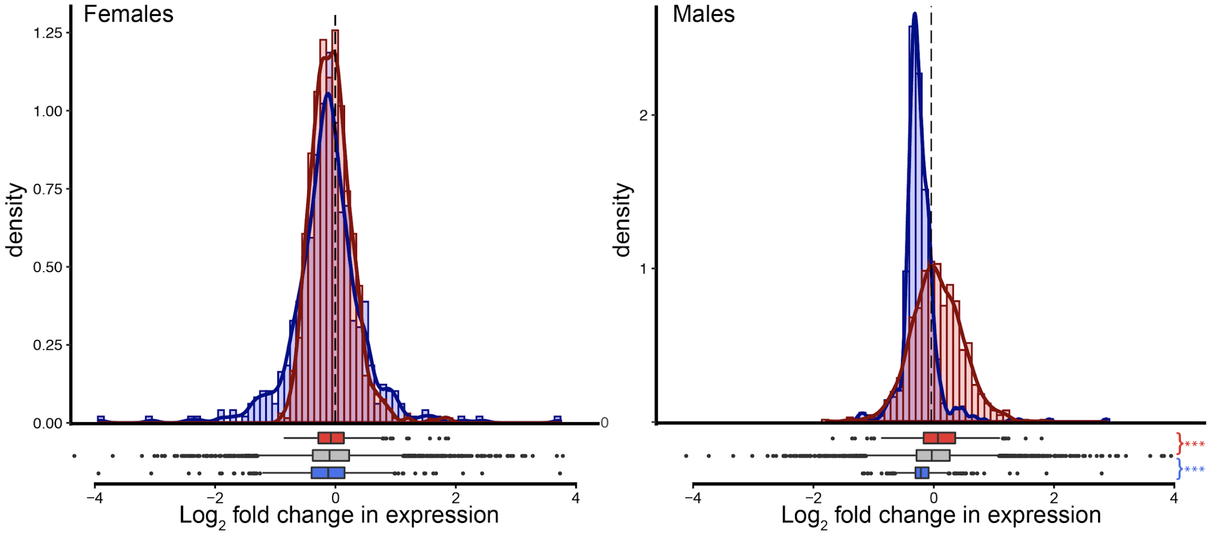


**Fig. S2 |** Expression shifts in genes sex-biased in both *D. montana* and *D. virilis* following cold treatment in females and males. Positive values indicate increased expression in cold-treated flies. Asterisks indicate the significance level (FDR) of Wilcoxon tests comparing the change in expression in female-biased (red) and male-biased (blue) genes to unbiased genes (***<0.001, **<0.01, *<0.05).

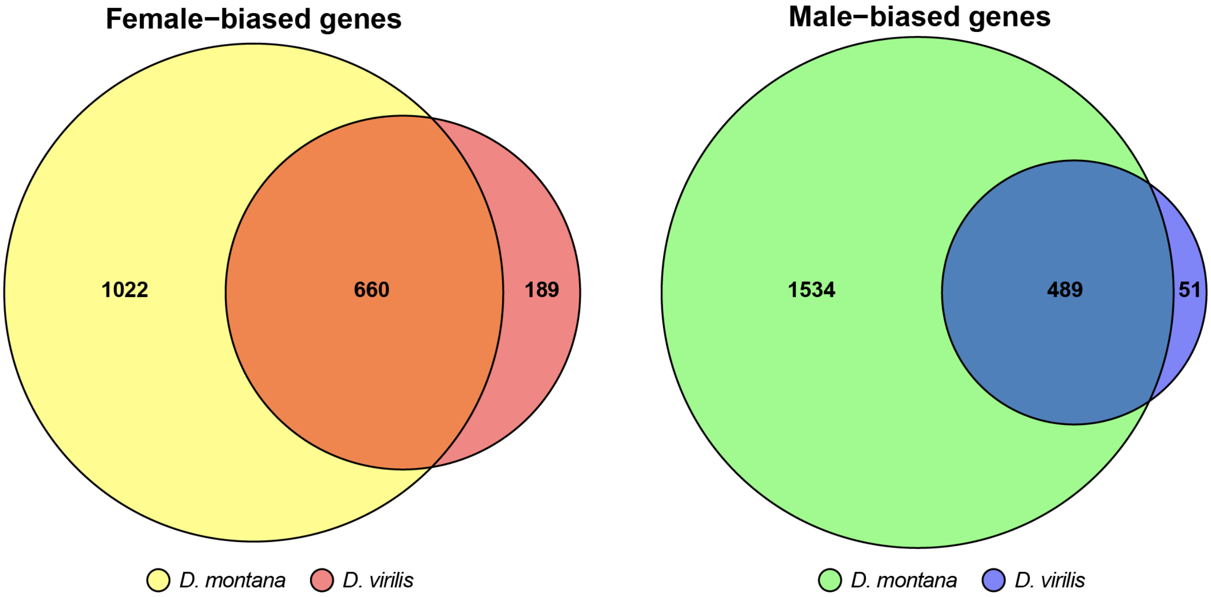


**Fig. S3 |** Overlap of sex-biased genes between *D. montana* and *D. virilis*

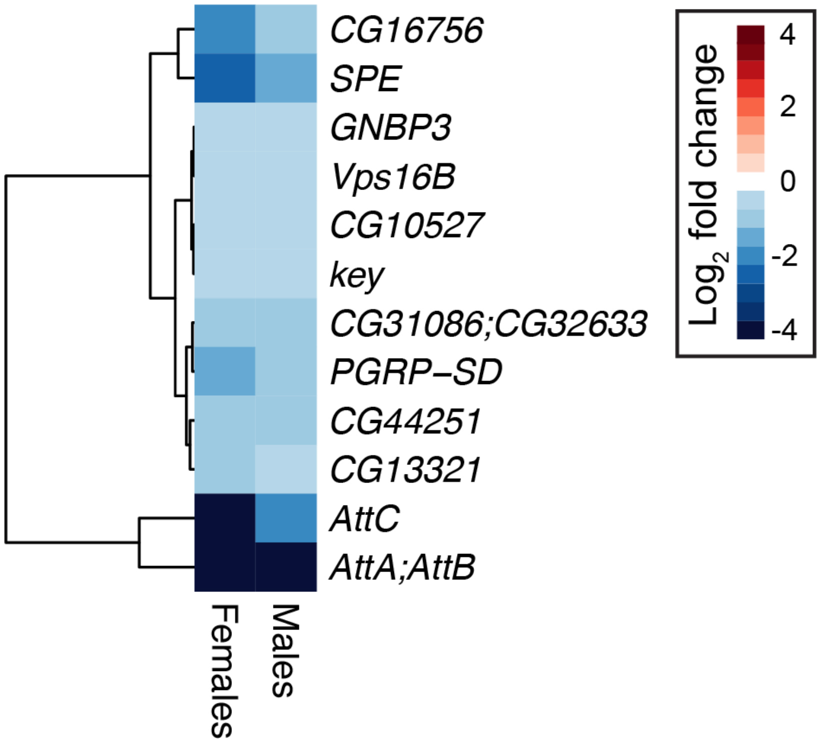


**Fig. S4 |** Expression shifts in genes enriched for innate immune response and DM9 repeats. Negative values indicate decreased expression in cold-treated flies.

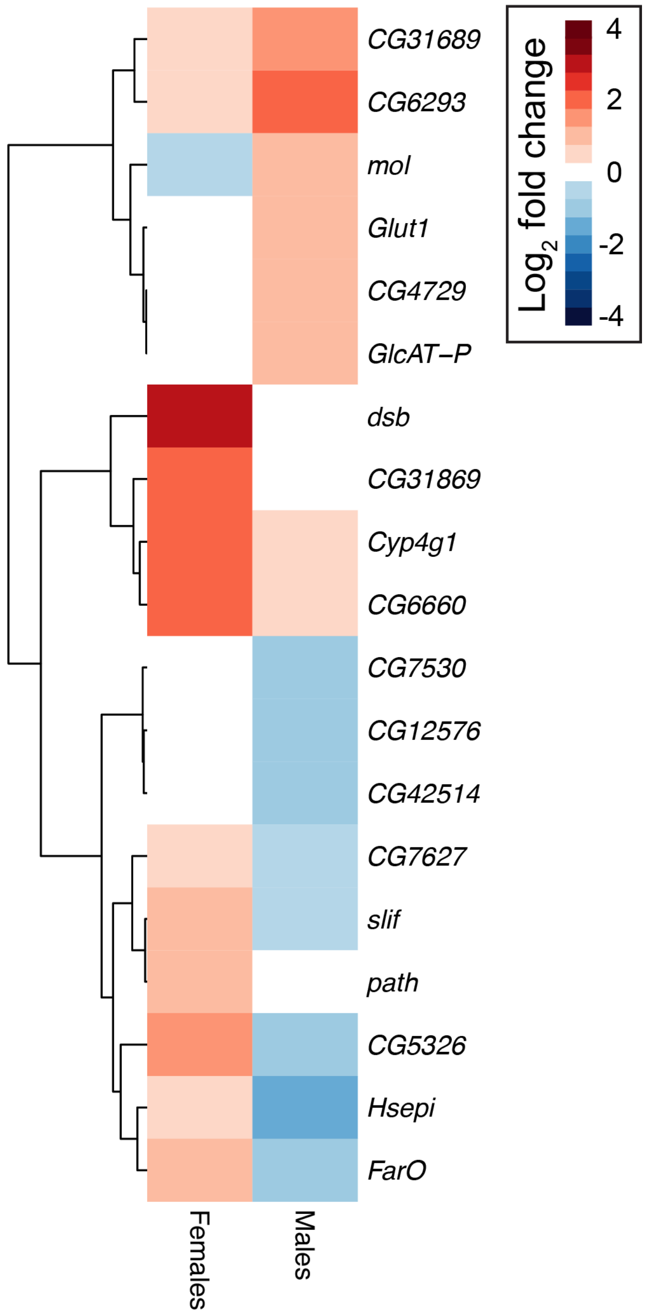


**Fig. S5 |** Expression shifts in genes with a significant sex by treatment interaction enriched for transmembrane transport, integral component of membrane processes. Positive values indicate increased expression in cold-treated flies.
